# A stainless-steel mortar, pestle and sleeve design for the efficient fragmentation of ancient bone

**DOI:** 10.1101/265587

**Authors:** Agata T Gondek, Sanne Boessenkool, Bastiaan Star

## Abstract

Ancient DNA (aDNA) bone extraction protocols routinely require the fragmentation of larger bone pieces into smaller pieces or powder prior to DNA extraction. To achieve this goal, different types of equipment such as oscillating ball mills, freezer mills, mortar and pestle, or drills can be used. Although all these approaches are suitable, practical drawbacks are associated with each method. Here, we present the design for a stainless-steel mortar and pestle, with a removable sleeve to contain bone material. The tool readily comes apart for ease of cleaning and its simplicity allows university workshops equipped with a lathe, boring tools and milling machine to make these components at local expense. We find that this design allows for the controlled fragmentation of ancient bone and that it significantly improves our sample throughput. We recommend this design as a useful, economical addition to existing laboratory equipment for the efficient handling of ancient bone.

## Methods Summary

We present an economical design of a stainless-steel mortar, pestle and sleeve for the efficient and controlled fragmentation of ancient bone.

To improve the economy and throughput of research projects, laboratory protocols require periodic evaluation with an aim to reduce human handling time and/or increase accuracy or efficiency. Such evaluation is of particular importance in ancient DNA (aDNA) laboratories where laboratory time associated with minimizing contamination forms a significant cost. Here, we present a design for an easily cleaned, stainless-steel mortar and pestle with a removable sleeve for fragmenting and pulverizing ancient bone, which is a fundamental procedure prior to DNA extraction (1). Even though a broad range of laboratory mills, grinders & crushers such as rotor-, knife-, disc- or ball-based mills, mortar grinders, jaw crushers or drills are available, not all tools can be easily adapted to clean room standards. Most commonly used equipment and approaches in aDNA studies are grinding balls with shaking or freezer mills (e.g. 2, 3-8), but also mortar and pestle (e.g. 9, 10, 11) or drills used at low rotational velocity (e.g. 12, 13). While these approaches are suitable, each method has practical disadvantages. For instance, the milling in oscillating ball or freezer mills occurs in a closed container and the process is therefore difficult to control in real time. This lack of control may lead to incompletely milled samples that need regularly inspection to ensure complete fragmentation or the over-processing of easily-fragmented samples. Moreover, these mills require time-consuming cleaning routines and represent a significant capital investment. While mortar and pestle are significantly cheaper and more straightforward to clean, it can be difficult to control impact and to keep fragments contained within the mortar, leading to the scattering of bone material throughout the working area. In addition, some samples may simply be too hard to be processed this way. Finally, the low rotational velocity required for drills to avoid the burning of bone results in substantial handling time in order to obtain sufficient powder. The design we present here addresses several of the drawbacks mentioned above and significantly improves our sample throughput.

The design consists of three parts *‒mortar, pestle and sleeve‒* that adhere closely together (Figure 1). The design is adapted from commercially available mortars that are used for the fragmentation of ores and minerals, for instance *Impact Mortar and Pestle* (cat. no 845/850 Chemplex®) or *Plattner’s mortar and pestle* (cat. no. 6883L10, Thomas Scientific) with several modifications for the specific purpose of fragmenting ancient bone. First, we elongated the pestle to provide easy grip and prevent fine bone powder from reaching hand level while exiting sleeve during repetitive grinding. Second, we elongated the sleeve to allow the pestle to remain in the sleeve to help contain the material. Three, we reduced the depth of the mortar chamber for easy access during cleaning. Fourth, we removed a one mm thick section over the entire length of the pestle. This allows the pestle to move more freely inside the mortar chamber and prevents the build-up of air-pressure. Initial tests of an earlier design without such section removed revealed that a tight fit of the pestle within the sleeve can result in the buildup of air-pressure with each downward move of the pestle. This pressure can then push fine bone powder through small seams between the mortar chamber and sleeve. Finally, each separate item (mortal, sleeve and pestle) is identified by an engraved number so that all pieces of a set can be kept together during handling.

**Figure 1.**
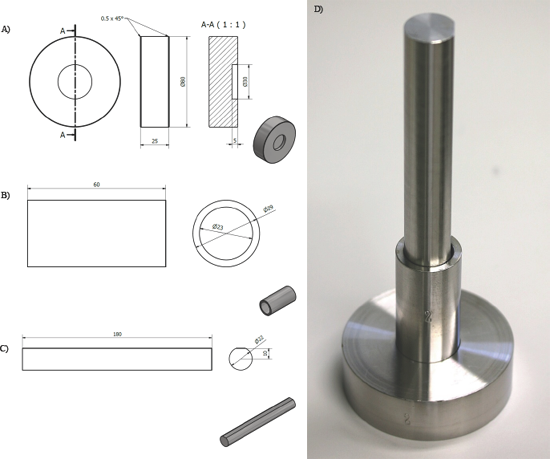
Design specifications of a mortar (A), sleeve (B) and pestle (C) for the fragmentation of ancient bone. All measurements are in millimetres. Each item is made of stainless-steel (AISI 316L). D) Photo of the assembled tool set.

All items have been constructed of stainless-steel (AISI 316L) in the Instrument Workshop at the University of Oslo using a lathe and boring tools. The mortar base is 25 mm high with a diameter of 80 mm, and the chamber has a depth of five mm with a diameter of 30 mm (Figure 1A). The sleeve has an outer diameter of 29 mm, inner diameter of 23 mm and a height of 60 mm (Figure 1B). The pestle has a diameter of 22 mm, a length of 180 mm and a one mm thick segment is removed from the transverse section over the entire length of the pestle (Figure 1C). All pieces are engraved with a respective set number (Figure 1D). Including labour (in Norway) and material expenses, the design costs approximate 150€ per set.

In our laboratory, the stainless-steel mortal, pestle and sleeve design presented here proved to be efficient for fragmenting and pulverizing a variety of bone samples, including specimens of fish and mammal. The material withstands a harsh cleaning routine and is corrosion resistant. The workflow presents no risk of overheating the bone and allows the realtime monitoring of the process of bone fragmentation. The tool proved to be especially useful while handling petrous bone samples, whereby the lower density parts of the bone break off preferentially after initial impacts of the pestle, making it easy to isolate the dense section of the petrous bone‒which contains most endogenous DNA (14, 15)‒ for further fragmentation. For particularly hard bone fragments, the tool is sufficiently robust to be used in combination with a rubber hammer.

We find that the three-piece stainless-steel mortar, pestle and sleeve is a versatile, economical and effective tool for fragmenting ancient bone. We have introduced specific modifications that are particularly useful for aDNA applications and find that the possibility to monitor the degree of bone pulverization while containing bone fragments within the sleeve provides a distinct advantage. The tool’s simple design allows university workshops equipped with a lathe, boring tools and a milling machine to make these sets at local expense, which ‒in our experience‒ is significantly more economical compared to commercially available varieties designed for ores and minerals. Moreover, the capital investment is substantially less than oscillating ball or freezer mills. We recommend this design as a useful, economical addition to existing laboratory equipment for the efficient handling of ancient bone.

## Author Contributions

BS & ATG conceptualized the design. ATG designed the mortal, pestle and sleeve. ATG tested all designs. BS and SB provided funding and consumables. BS & ATG wrote the manuscript in collaboration with SB.

## Acknowledgements

We thank Jan Kristiansen from University of Oslo Instrumental Workshop for help with designing and manufacturing the mortar, pestle and sleeve. This work was supported by the Research Council of Norway (grant numbers 262777 to BS and 230821 to SB).

## References

1. Morales Colón E, Hernández M, Candelario M, Meléndez M, & Dawson Cruz T 2017. Evaluation of a Freezer Mill for Bone Pulverization prior to DNA Extraction: An Improved Workflow for STR Analysis. Journal of forensic sciences.

2. Rohland N & Hofreiter M 2007. Ancient DNA extraction from bones and teeth. Nature Protocols 2:1756.

3. Mitchell KJ, Llamas B, Soubrier J, Rawlence NJ, Worthy TH, Wood J, Lee MSY, & Cooper A 2014. Ancient DNA reveals elephant birds and kiwi are sister taxa and clarifies ratite bird evolution. Science 344:898-900.

4. Boessenkool S, Hanghøj K, Nistelberger HM, Der Sarkissian C, Gondek A, Orlando L, Barrett JH, & Star B 2016. Combining bleach and mild pre - digestion improves ancient DNA recovery from bones. Molecular Ecology Resources.

5. Lazaridis I, Nadel D, Rollefson G, Merrett DC, Rohland N, Mallick S, Fernandes D, Novak M, et al 2016. Genomic insights into the origin of farming in the ancient Near East. Nature 536:419.

6. Palacio P, Berthonaud V, Guérin C, Lambourdière J, Maksud F, Philippe M, Plaire D, Stafford T, et al 2017. Genome data on the extinct Bison schoetensacki establish it as a sister species of the extant European bison (Bison bonasus). BMC Evol. Biol. 171:48.

7. Star B, Boessenkool S, Gondek AT, Nikulina EA, Hufthammer AK, Pampoulie C, Knutsen H, André C, et al 2017. Ancient DNA reveals the Arctic origin of Viking Age cod from Haithabu, Germany. Proceedings of the National Academy of Sciences 11434:9152-9157.

8. Kennett DJ, Plog S, George RJ, Culleton BJ, Watson AS, Skoglund P, Rohland N, Mallick S, et al 2017. Archaeogenomic evidence reveals prehistoric matrilineal dynasty. Nature Communications 8:14115.

9. Knapp M, Clarke AC, Horsburgh KA, & Matisoo-Smith EA 2012. Setting the stage-building and working in an ancient DNA laboratory. Annals of Anatomy- Anatomischer Anzeiger 1941:3-6.

10. Fortes GG, Grandal-d’Anglade A, Kolbe B, Fernandes D, Meleg IN, García-Vázquez A, Pinto-Llona AC, Constantin S, et al 2016. Ancient DNA reveals differences in behaviour and sociality between brown bears and extinct cave bears. Mol Ecol 2519:4907-4918.

11. Matisoo-Smith EA, Gosling AL, Boocock J, Kardailsky O, Kurumilian Y, Roudesli-Chebbi S, Badre L, Morel J-P, et al 2016. A European Mitochondrial Haplotype Identified in Ancient Phoenician Remains from Carthage, North Africa. PloS one 115:e0155046.

12. Adler C, Haak W, Donlon D, Cooper A, & Consortium G 2011. Survival and recovery of DNA from ancient teeth and bones. Journal of Archaeological Science 385:956-964.

13. Sirak K, Novak M, & Cheronet O 2017. A minimally-invasive method for sampling human petrous bones from the cranial base for ancient DNA analysis. BioTechniques 626:283-289.

14. Pinhasi R, Fernandes D, Sirak K, Novak M, Connell S, Alpaslan-Roodenberg S, Gerritsen F, Moiseyev V, et al 2015. Optimal Ancient DNA Yields from the Inner Ear Part of the Human Petrous Bone. PLOS ONE 106:e0129102.

15. Hansen HB, Damgaard PB, Margaryan A, Stenderup J, Lynnerup N, Willerslev E, & Allentoft ME 2017. Comparing Ancient DNA Preservation in Petrous Bone and Tooth Cementum. PLOS ONE 121:e0170940.

